# Synthesis and Characterization of Phase-Separated Extracellular Condensates in Interactions with Cells

**DOI:** 10.1101/2025.03.24.644961

**Authors:** Aida Naghilou, Tom M.J. Evers, Oskar Armbruster, Vahid Satarifard, Alireza Mashaghi

**Affiliations:** Medical Systems Biophysics and Bioengineering, Leiden Academic Centre for Drug Research, Faculty of Science, Leiden University, 2333CC, Leiden, The Netherlands; Laboratory for Interdisciplinary Medical Innovations, Centre for Interdisciplinary Genome Research, 2333CC, Leiden University, Leiden, The Netherlands; Institute of Synthetic Bioarchitectures, Department of Biotechnology and Food Science, BOKU University, 1190 Vienna, Austria; Yale Institute for Network Science, Yale University, New Haven, CT 06520, USA

## Abstract

Biomolecular condensates formed through liquid-liquid phase separation play key roles in intracellular organization and signaling, yet their function in extracellular environments remains largely unexplored. Here, we establish a model using heparan sulfate, a key component of the extracellular matrix, to study extracellular condensate-cell interactions. We demonstrate that heparan sulfate can form condensates with a positively charged counterpart in serum-containing solutions, mimicking the complexity of extracellular fluid, and supporting cell viability. We observe that these condensates adhere to cell membranes and remain stable, enabling a versatile platform for examining extracellular condensate dynamics and quantifying their rheological properties as well as their adhesion forces with cellular surfaces. Our findings and methodology open new avenues for understanding the organizational roles of condensates beyond cellular boundaries.

## Introduction

Phase-separated biomolecular condensates have emerged as key players in cellular compartmentalization, enabling a range of critical functions such as gene regulation, membrane remodeling, and intracellular sensing ^1-6^. Liquid-liquid phase separation has been identified as the underlying mechanism of condensate formation and has been linked to physicochemical properties of biomolecules^7-9^. Although much of the research on biomolecular condensates has focused on their roles inside cells, recent findings suggest that extracellular molecules can also form condensates ^10-12^. These observations raise questions about the role of condensates in extracellular organization, where they may contribute to interactions with cells and the extracellular matrix. Despite these possibilities, data on extracellular condensates and their interactions with cells are scarce. Developing assays for studying extracellular condensates and their cellular interactions could open new avenues for understanding and engineering biochemical processes, as phase separation may modulate enzyme kinetics, transport processes, and mechanochemistry ^13,14^.

Modeling condensate-cell interactions in an extracellular setting requires considerations distinct from those for intracellular condensates. While many studies use simplified buffer systems to mimic the intracellular environment by introducing ions and crowding agents ^15-18^, extracellular condensates require assays in culture medium conditions that support cell viability. In addition, the extracellular fluid in humans resembles plasma ^19^, implying that extracellular condensates would encounter plasma proteins and other soluble factors, necessitating the development of a condensate model that sustains stability within such an environment to study its properties.

One characteristic feature of the extracellular environment is its abundance of charged polysaccharides, which can contribute to condensate formation alongside proteins. Glycosaminoglycans (GAGs) are long, unbranched, and highly negatively charged polysaccharides that are key components of the extracellular environment ^20^. They are involved in a variety of biological and pathological processes, including development, angiogenesis, and inflammatory responses ^21,22^. One such GAG, heparan sulfate (HS), interacts with a broad spectrum of molecules at the cellular interface, playing a vital role in extracellular signaling ^23-25^. Heparin (H) is another GAG known for its anticoagulant properties ^26^ has been shown to form coacervates with charged proteins ^23,24^. Furthermore, *in silico* approaches have predicted the condensate formation propensity of numerous extracellular proteins ^27^. These lines of evidence suggest that phase separation of GAGs and extracellular proteins is ubiquitous and may play a role in various (patho)physiological settings.

In this article, we demonstrate that HS phase separates to form condensate droplets in the presence of a positively charged counterpart. Importantly, these condensates are also formed in serum-like media containing proteins and nutrients necessary to sustain cell viability. We used a combination of optical and scanning probe microscopy (SPM) as well as fluorescence recovery after photobleaching (FRAP) to investigate the liquid-like behavior of these condensates. In addition, we developed assays enabling the determination of condensate mechanical properties such as viscoelasticity and adhesion forces on cells, marking the first application of SPM in analyzing cell-condensate interactions. Our assay is independent of condensate nature and cell types, providing a versatile and adaptable platform for investigating interactions between extracellular condensates, cells, and the extracellular matrix (ECM).

## Results

### Heparan sulfate phase separates with poly-L-lysine with distinct differences to heparin

While H phase separates with poly-L-lysine (pK) at 1 M KCl with crowding agents, no droplets were formed with pK-HS at the same buffer concentration (Figure 1A). To gain insights into the nature of condensation of pK-HS, we performed a concentration gradient analysis of KCl to determine the conditions at which condensate formation occurs (Figure 1B). At lower salt contents, pK-HS phase separation leads to condensate formation, whereas at higher KCl concentrations, the system transitions back to a single-phase regime, highlighting the differences between HS and H condensates. Notably, pK-H interactions at low salt concentrations, comparable to those that leads to pK-HS droplet formation, causes an aggregation of pK-H, as depicted in Figure 1C. As 0.15 M KCl is routinely used for studying biological systems, for further investigation of pK-HS condensates we employed this salt concentration.

**Figure 1:**
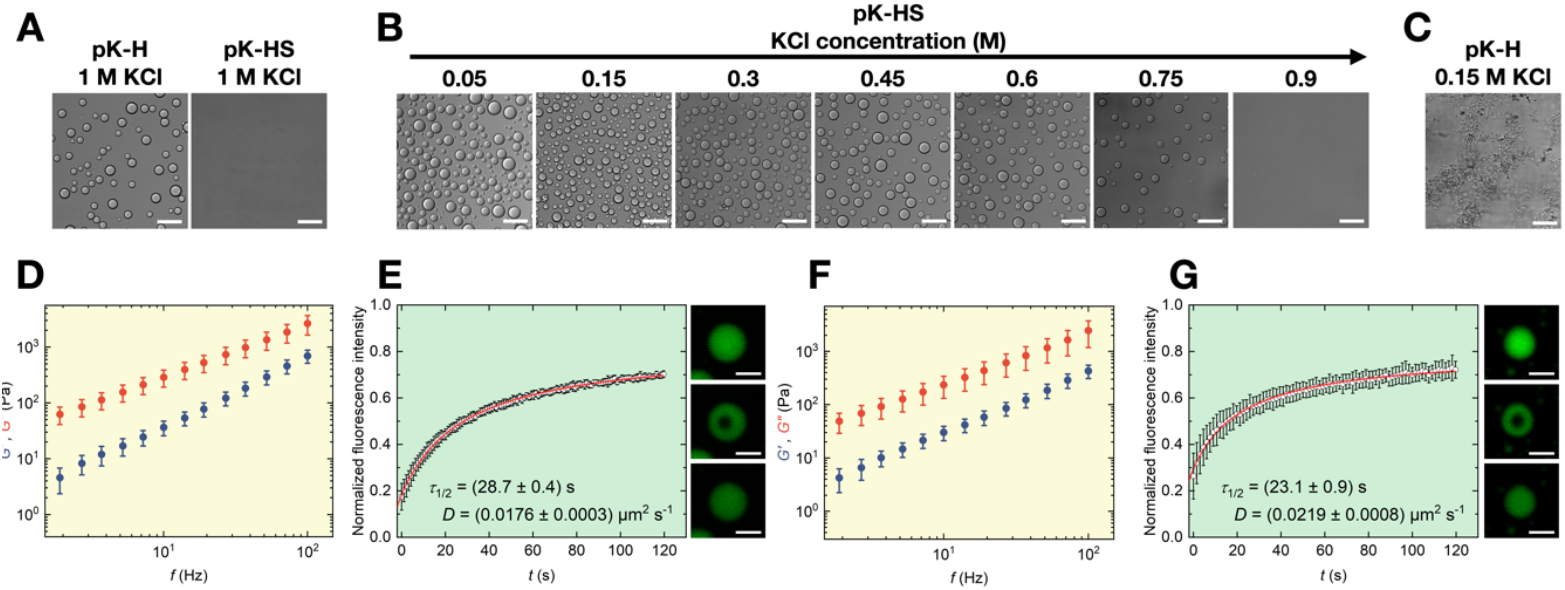
pK-HS condensates differ from pK-H. **A)** Representative confocal micrographs of pK-H droplets and pK-HS at 1 M KCl, scale bar is 50 µm. **B)** Exemplary confocal micrographs of pK-HS droplets at various KCl concentrations, scale bar is 50 µm. **C)** Representative confocal micrographs of pK-H droplets at 0.15 M KCl **D)** Elastic (*G*’, blue) and viscous (*G*”, red) shear moduli of pK-HS condensates formed at 0.15 M KCl and RT, measured at RT (mean ± SD, n = 3). **E)** Time-dependent normalized fluorescence intensity depicting the recovery of the bleached area (circular data points) and the fit with Eq. 2, resulting in half-time of recovery (τ_1/2_) and diffusion constant (*D*) with Eq. 3 pK-HS condensates formed at 0.15 M KCl and RT, measured at RT (mean ± SD, n = 1) as well as representative confocal micrographs of droplet recovery after photobleaching, scale bar is 10 µm. **F)** Elastic (*G*’, blue) and viscous (*G*”, red) shear moduli of pK-HS condensates formed at 0.15 M KCl and RT, measured at 37°C (mean ± SD, n = 3). **G)** Time-dependent normalized fluorescence intensity depicting the recovery of the bleached area (circular data points) and the fit with Eq. 2, resulting in half-time of recovery (τ_1/2_) and diffusion constant (*D*) with Eq. 3 for pK-HS condensates formed at 0.15 M KCl and RT, measured at 37°C (mean ± SD, n = 1) as well as representative confocal micrographs of droplet recovery after photobleaching, scale bar is 10 µm.

In order to demonstrate the liquid-like nature of pK-HS condensates and to decipher their viscoelastic characteristics, we employed SPM and FRAP. Figure 1D displays the rheological properties of droplets at room temperature (RT). The real (elastic modulus, *G*’) and imaginary (viscous modulus, *G*”) shear moduli are presented as blue and red circles, respectively. The larger values of *G*” in comparison to *G*’ demonstrated a dominant viscous behavior of droplets at all experimentally accessible frequencies. A linear fit of *G*” was performed to determine the viscosity, η which was found to be (26.1 ± 0.5) Pa (**Error! Reference source not found**.A). The surface energy density of the droplets, γ was calculated from the quasi-static indentation curve (see Methods) and is shown in in **Error! Reference source not found**.B. Average recovery traces, the half-time of recovery, τ_1/2_ and diffusion coefficients, *D* as well as representative confocal images of condensates during FRAP experiments is exemplified in Figure 1E, also indicating the liquid-like behavior of pK-HS condensates based on the fast recovery.

As we aimed to study cell-condensate interactions, a crucial prelude is biological temperatures. Hence, we repeated the SPM and FRAP measurements after heating the droplets to 37°C, shown in Figure 1F and G, respectively. In accordance with Arrhenius’ law ^28^, η decreased at 37°C to (23.5 ± 0.7) Pa s. This is also indicated by the smaller values of τ_1/2_ and larger *D* at higher temperature. A comparison of the material properties of condensates at RT and 37°C is depicted in **Error! Reference source not found**..

### Heparan sulfate phase separates with poly-L-lysine in a serum-containing medium without the need for crowding agents or surface passivation

Deciphering condensate-cell interactions in the extracellular environment requires culture conditions that preserve cell viability. This prompted us to study whether pK-HS condensates also form in a culture medium. Furthermore, extracellular fluid in human tissues is rich in plasma proteins, which may impact condensate formation and properties. As such, we prepared serum containing media typically used for *in vitro* cell culture studies (see Methods). pK and HS were added to the medium with the same end concentrations used in buffer. Figure 2 A shows phase contrast micrographs of the pK-HS condensates in the culture medium, with morphologies similar to those formed in buffer. To confirm that the condensates are indeed pK-HS and not the result of interactions between any medium components with either pK or HS, we repeated the experiments using medium with added pK without HS (Figure 2 B top), and vice versa (Figure 2 B, bottom). No condensates were formed unless both pK and HS were present, thereby demonstrating that the droplets formed are indeed pK-HS condensates. Interestingly, due to the high protein content of culture medium, neither a crowding agent nor a surface coating was necessary. Condensates retained their natural spherical morphology, despite a lack of any surface passivation, which is in contrast to the pK-HS condensates prepared in buffer (**Error! Reference source not found**.). This hints at the fact that the molecules of the culture medium may have bounded to the surface of the dish, acting as a passivation. To gain deeper insights on the differences between H and HS, we also investigated pK-H interactions in the same culture media as used for pK-HS. Consistent with the behavior seen in buffer (Figure 1A and C), while pK-HS leads to condensate formation, pK-H forms aggregates in culture medium (**Error! Reference source not found**.)

**Figure 2:**
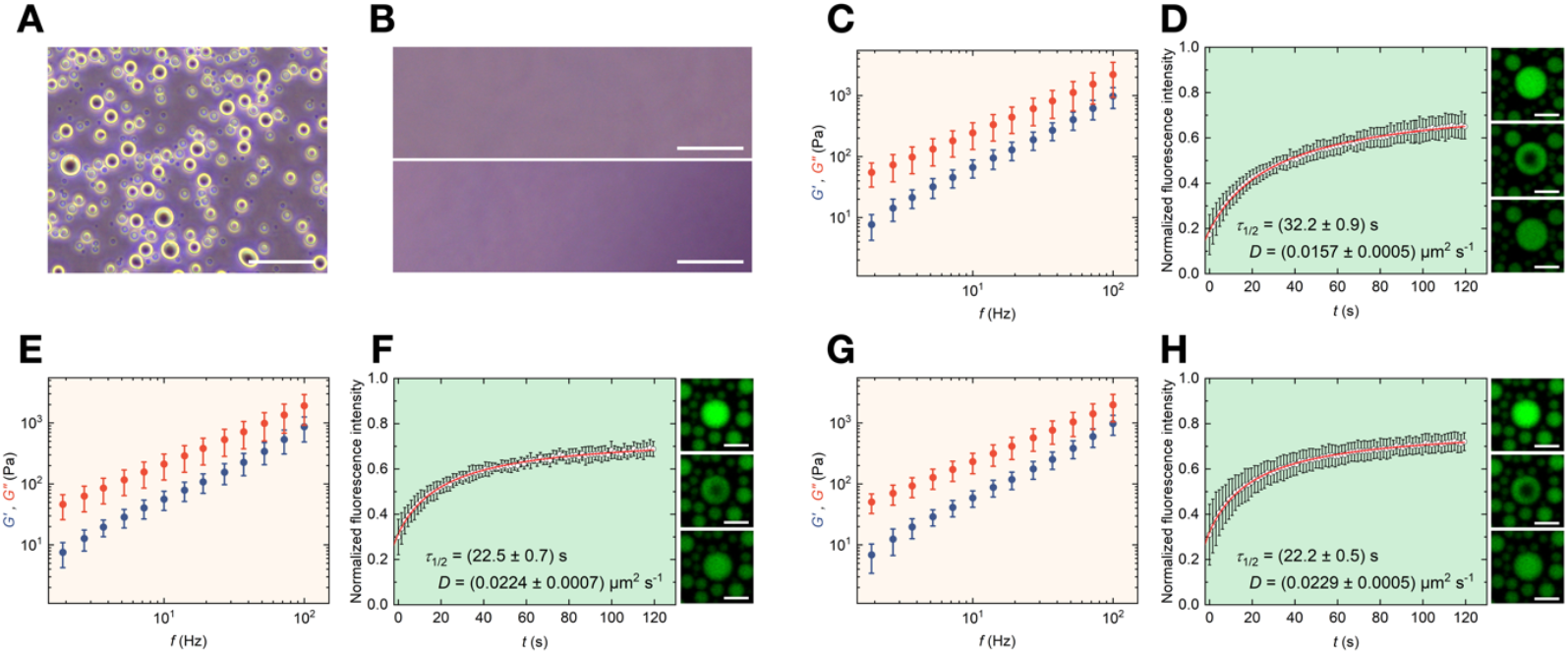
pK-HS condensates formed in culture medium. **A)** Representative phase contrast micrograph of pK-HS droplets formed in culture medium; scale bar is 50 µm. **B)** Exemplary phase contrast micrographs of culture medium with added pK without HS (top) and with added HS without pK (bottom), scale bar is 50 µm. **C)** Elastic (*G*’, blue) and viscous (*G*”, red) shear moduli of pK-HS condensates formed in culture medium and RT, measured at RT (mean ± SD, n = 3). **D)** Time-dependent normalized fluorescence intensity depicting the recovery of the bleached area (circular data points) and the fit with Eq. 2, resulting in half-time of recovery (τ_1/2_) and diffusion constant (*D*) with Eq. 3 pK-HS condensates formed in culture medium and RT, measured at RT (mean ± SD, n = 1) as well as representative confocal micrographs of droplet recovery after photobleaching, scale bar is 10 µm. **E)** Elastic (*G*’, blue) and viscous (*G*”, red) shear moduli of pK-HS condensates formed in culture medium and RT, measured at 37°C (mean ± SD, n = 3). **F)** Time-dependent normalized fluorescence intensity depicting the recovery of the bleached area (circular data points) and the fit with Eq. 2, resulting in half-time of recovery (τ_1/2_) and diffusion constant (*D*) with Eq. 3 pK-HS condensates formed in culture medium and RT, measured at 37°C (mean ± SD, n = 1) as well as representative confocal micrographs of droplet recovery after photobleaching, scale bar is 10 µm. **G)** Elastic (*G*’, blue) and viscous (*G*”, red) shear moduli of pK-HS condensates formed in culture medium and 37°C, measured at 37°C (mean ± SD, n = 3). **H)** Time-dependent normalized fluorescence intensity depicting the recovery of the bleached area (circular data points) and the fit with Eq. 2, resulting in half-time of recovery (τ_1/2_) and diffusion constant (*D*) with Eq. 3 pK-HS condensates formed in culture medium and 37°C, measured at 37°C (mean ± SD, n = 1) as well as representative confocal micrographs of droplet recovery after photobleaching, scale bar is 10 µm.

The material properties of pK-HS condensates in medium vary in comparison to those formed in buffer. SPM measurements (Figure 2C) showed that, the condensates are closer to their cross-over point at higher frequencies and their η at RT is slightly lower (21.7 ± 0.7) Pa s than the pK-HS droplets in buffer. Interestingly, we did not see the same trend in FRAP (Figure 2D), where τ_1/2_ was in fact slightly higher than the values for measurements in buffer at RT. In contrast to the slight changes of η between medium and buffer, γ values were noticeably lower for pK-HS droplets formed in medium in comparison to those in buffer (**Error! Reference source not found**.B). This behavior could be related to the absent of the crowding agent in the medium, greatly impacting the interfacial γ. Afterwards, we increased the temperature from RT to 37°C, and observed lower viscosities in SPM (Figure 2E), as well as smaller τ_1/2_ and larger *D* in FRAP (Figure 2F).

For studying condensate interactions with cells, it is crucial to avoid temperature fluctuations for cells. Hence, we also investigated the phase separation with medium pre-heated to 37°C before adding pK and HS to form condensates. The resulting droplets showed material properties comparable to those formed by heating the medium from RT to 37°C, with lower η and faster fluorescent recovery in comparison to pk-HS condensate prepared at RT, as shown in Figure 2G and H. A comparison of material properties of droplets in all conditions measured by SPM and FRAP is depicted in **Error! Reference source not found**.. These results demonstrate that pK-HS droplets serve as a suitable model system for studying condensate-cell interactions, as they form under biologically relevant fluids and temperature while retaining their liquid-like properties.

### Scanning probe microscopy reveals cell-condensate interactions

After demonstrating that pK-HS condensates remain stable in culture medium and maintain their liquid-like characteristics, we proceeded to investigate their interactions with fibroblast cells (Figure 3A). For this purpose, pK and HS were mixed to a fresh culture medium at 37°C and immediately added on the cells. An exemplary phase contrast micrograph after 2 h of incubation, depicting the condensates settling both on top and next to the cells is shown in Figure 3B. The condensate size on the cells is larger in comparison to the ones on the dish, which may be due to the movement of cells, increasing the chances of condensate fusion. This is an indication that cells are viable and migrating within the pK-HS containing medium, also visible by similar morphologies of cells in culture medium without and with condensates (Figure 3A and B, respectively).

**Figure 3:**
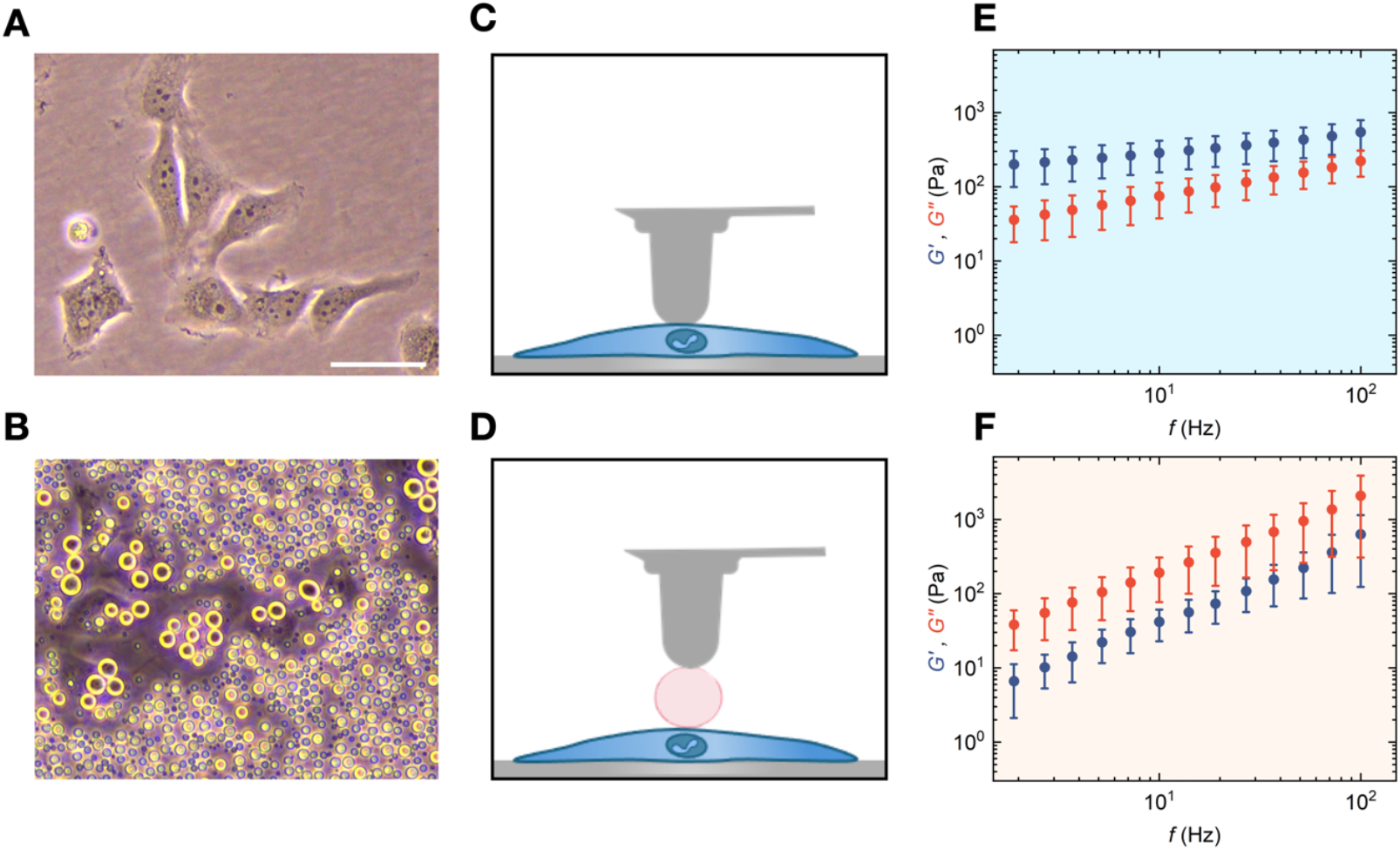
pK-HS condensates’ rheological properties in contact with fibroblast cells. **A)** Representative phase contrast micrograph of fibroblast cells in culture medium, scale bar is 50 µm. **B)** Exemplary phase contrast micrographs of pK-HS condensates in culture medium residing on fibroblast cells, scale bar is 50 µm. **C)** Schematic representation of the SPM probe, indenting the fibroblast cells. **D)** Schematic representation of the SPM probe, indenting the condensates residing on cells **D)** Fibroblast cells in culture medium at 37°C **E)** Elastic (*G*’, blue) and viscous (*G*”, red) shear moduli of fibroblast cells measured at 37°C (mean ± SD, n = 4). **F)** Elastic (*G*’, blue) and viscous (*G*”, red) shear moduli of pK-HS condensates on the fibroblast cells formed in culture medium and 37°C, measured at 37°C (mean ± SD, n = 3).

To determine the mechanical properties of droplets resting on cells, it is crucial to consider that the cells’ elastic moduli are much lower than those of polystyrene dishes, which may lead to a deformation of cell while indenting the condensate. To address this, we derived a stacked Hertzian contact model, allowing us to extract the *G*’ and *G*” of condensates by taking into account the shear moduli of the cell as a known parameter (see Methods). Hence, we first measured the rheological properties of the cells and in the same manner the condensates resting on cells (Figure 3C and D, respectively). Both *G*’ and *G*” of fibroblast cells show a rise at higher frequencies with a dominant *G*’, hinting at a mainly elastic behavior (Figure 3E). Employing the stacked Hertzian contact model (Eq. 4-10), the viscoelastic characteristics of condensates on cells were determined (Figure 3F). The condensates on cells showed similar η and slightly higher γ in comparison to those formed in medium without the presence of cells. A comparison of η and γ for condensates in medium, with and without the cells is depicted in Supplementary Figure S4.

Next, we measured the adhesion forces between the condensates and the cells with SPM (Figure 4A). To achieve this, a tipless cantilever was coated with laminin, chosen due to its strong interactions with HS ^29^. A representative force (*F*) curve as a function of time (*t*) is shown in Figure 4B, illustrating the measurement process. During the experiment, the cantilever head was lowered towards the condensate until a predefined deflection was achieved (indicated by the green line in Figure 4B). This was followed by a pause period (red line) to allow sufficient time for the laminin-coated cantilever to bind firmly to the condensate. Subsequently, the cantilever was retracted, initiating the detachment phase of the curve, represented by the dark grey line.

**Figure 4:**
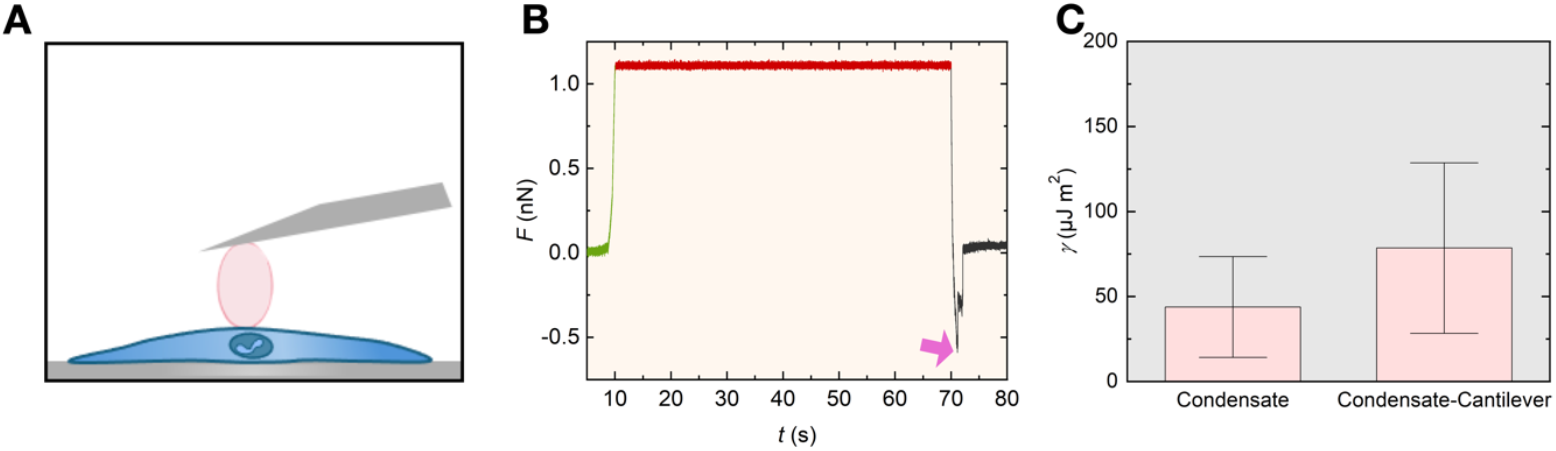
pK-HS condensates’ adhesion forces with fibroblast cells. **A)** Schematic representation of the SPM cantilever, detaching the condensates residing on cells. **B)** A representative curve of cantilever deflection force over time, *F(t)* showing the approach in green, pause in red, and retraction in dark grey. The violet arrow indicates the lowest force value during detachment, corresponding to the adhesion force *F*_ad_. **C)** Comparison of the interfacial surface energy density (γ) between the pK-HS condensates condensate and dilute phase denoted as condensate and between the condensate and laminin coated cantilever denoted as condensate-cantilever on the fibroblast cells formed in culture medium and 37°C, measured at 37°C (mean ± SD, n = 3).

Importantly, the alignment of the baseline during the approach and detachment phases demonstrates that while the cantilever lost contact with the condensate, the condensate itself remained securely attached to the cell surface. This observation underscores the strong adhesive interaction between the condensates and the cellular surface, hinting that the pK-HS droplets could function as a biological adhesive on the cell. The minimum force value observed during the detachment phase corresponds to the adhesion forces *F*_*ad*_ between the cantilever and the cell (highlighted by the violet arrow) and averages to 0.98 ± 0.09 nN. The distribution of the adhesion forces for all measurements is depicted in Supplementary Figure S 5.

Employing the JKR model, and droplet sizes (see Methods, Eq. 11) the interfacial surface energy distribution γ between the droplet and the cantilever were calculated. This was compared with γ determined from the quasi-static indentation curves while considering the contribution of the cell deformation to the overall indentation (See Methods, Eq. 12-15). The results of the comparison are shown in Figure 4C. The higher γ between the condensate and the cantilever in comparison to the condensate and dilute phase indicates the adhesive interactions of the pK-HS droplets with the laminin coated cantilever.

## Discussion and Conclusions

Liquid-liquid phase separation underpins key cellular functions ^30^. It drives the formation of functional condensates in the cytoplasm and nucleoplasm ^31,32^ and can occur on the plasma membrane or endoplasmic reticulum, supporting condensate formation that aids in cell signaling, tight junctions, and synaptic transmission ^33,34^. The intracellular environment with its high protein density facilitates liquid-liquid phase separation ^23^, and thus has been the primary focus on condensate research over the last decade. The extracellular environment of tissues is also rich in biopolymers with a predicted propensity for condensate formation ^27^, and recent studies have experimentally identified the presence of molecules in the extracellular environment capable of forming condensates, including elastin and galectin-3 ^10,11^. The discovery of extracellular condensates is paramount as the condensates may reside on cells, a configuration that could enable various mechanical, biochemical, and transport processes at the cellular interface ^23^. These findings underscore the need for technical development that enable characterization of extracellular condensates and their cellular interactions to understand their roles and mechanisms. In this study, we developed a methodology using SPM for quantifying interactions between extracellular condensates and cells. While SPM has recently been utilized to study biomolecular condensates, particularly for probing liquid-gel phase transitions ^35^, our work extends its potential for analyzing extracellular condensates and establishes the groundwork for precise measurements of their interactions with cellular surfaces.

In this regard, we created biomolecular condensates in an extracellular fluid mimic. Widely present on cell surfaces, within the ECM, basement membranes, and even intracellularly, HS plays a pivotal role in numerous biological processes ^36^. Through dynamic multivalent ionic interactions, it binds various positively charged extracellular proteins, aiding in the structural organization of the ECM ^25^. Furthermore, its interactions with positively charged molecules, such as chemokines and growth factors, have been shown to results in condensate formation ^23,24^. Given its crucial role in the ECM and its affinity for positive molecules, we utilized pK-HS condensates in a culture medium containing components that support cell viability. These condensates exhibited morphologies similar to those formed in the buffer system with crowding agents. However, in the culture medium, neither additional crowding agents nor surface passivation were needed, suggesting that the protein-rich environment provided by serum acts as a natural crowding agent, while protein-surface interactions mimic surface passivation. We characterized the material properties of pK-HS condensates using SPM and FRAP at room and physiological temperatures. Our results demonstrated that condensate viscosity decreases at higher temperatures, which is in line with previous studies investigating the impact of temperature on condensates ^28^. While H is often used as a mimetic for HS in biochemical interaction studies, our study reveals clear differences in condensation behaviors of HS and H. Specifically, under the same conditions in which HS forms liquid condensates, H instead forms aggregates. As such, we recommend that for extracellular condensate studies, H should not be used as a replacement for HS. These distinct behaviors and physicochemical properties may be explained the differing roles of HS and H in the extracellular space ^37^.

We observed that pK-HS condensates form in culture media containing cells and stably adhere to cellular surfaces. Previous studies have highlighted the potential role of condensates in mediating cell-cell adhesion, suggesting that they may act as molecular bridges that facilitate intercellular connections ^10,38^. Our findings not only align with these observations but also provide a new dimension by enabling the quantitative assessment of condensate properties during cell interactions. Using SPM and introducing a novel stacked Hertzian model, we were able to measure the rheological properties of these condensates while they were directly on the cells. In addition to rheology, by determination of surface tension, as well as adhesion forces with SPM, we offer a novel approach to understanding the mechanical dynamics of condensates on cells, paving the way for deeper insights into their biological implications, including their roles in cellular communication and tissue organization.

Condensate interaction with neighboring cells is fundamentally important with many implications for biology and therapeutic development. These direct physical interactions could enable various mechanical, biochemical, and transport processes at the condensate-cell interface. By organizing ECM components, receptor ligands, or signaling molecules near the cell surface, these condensates may influence cell behavior through biochemical and mechanical processes^21^. They could enhance or inhibit enzymatic reactions, regulate substrate transport, and provide spatial control over signaling pathways ^22^. Acting as dynamic microenvironments, extracellular condensates may regulate the availability and diffusion of key molecular players, thereby potentially exerting local control over nutrient uptake, drug delivery, or growth factor responses ^39^. Additionally, their adhesion to cell membranes suggests a role in modulating cellular mechanical properties, such as stiffness or elasticity, with implications for processes like migration, mechanosensing, and ECM remodeling ^12^. Condensates’ role in regulating mechanics could be particularly relevant in pathological contexts, such as cancer, where ECM organization is disrupted ^40^. Future investigations will shed light on how extracellular condensates mediate interactions with the cells, the surrounding matrix, and potentially contribute to ECM remodeling.

## Methods

### Condensate preparation

Experiments were performed on virgin polystyrene culture dishes (93040, TPP, Switzerland), with reagents purchased from Sigma-Aldrich unless noted otherwise. Dishes were treated with 1%w/v BSA (A2153) for 30 minutes followed by rinsing 5–6 times with deionized water. For pK-H condensates, the coacervation buffer contained 1 M KCl (p9333), 10 mM imidazole (56750), 1 g/l NaN3 (8.22335), and 50 g/l Ficoll® PM 70 (F2878). H (H3393) and pK (P2658) were added sequentially with 30 µM concentration for each. For the pK-HS condensates, the same buffer was used except for the concentration of KCl, which was varied between 0.05 and 1 M to determine the best condition, which was chosen as 0.15 M KCl, for all other pK-HS experiments. Same pK and HS (H7640) concentrations of 30 µM were employed.

For pK-HS droplet experiments in culture medium, DMEM (Capricorn, Germany) containing glucose (4.5 g/l) and Sodium Pyruvate was supplemented with 10% fetal calf serum, 2.5% HEPES, and 1% penicillin-streptomycin. Afterwards without any further additives, pK and HS were added to an end concentration of 30 µM each.

### Imaging of droplets and cells

For imaging the morphology of condensates as shown in Figure 1, they were prepared in an ultra-low binding 384-Wells (Corning™, USA) and confocal microscopy was carried out with a Nikon Ti2 Eclipse microscope and a 20x/0.75 objective. Phase contrast microscopy as seen in Figure 2 and 3 was performed with a Nikon Ti2 microscope and a 20x/0.4 objective and the droplets and cells were prepared in uncoated polystyrene TPP culture dishes.

### Fluorescent Recovery After Photobleaching (FRAP)

For FRAP, a 5% FITC-labeled pK (P3543) solution was mixed with the unlabeled counterpart. Experiments were conducted using a Nikon Ti2 Eclipse microscope with a 20x/0.75 objective. With a 488 nm laser and an average power of 250 µW, a region with radius of *r*n = 1.5 µm on the droplets was bleached for 1 s, followed by a 2 min recovery scan. This procedure was repeated for 7-9 condensates in one sample preparation. Data were normalized to an unbleached reference droplet to correct for photodecay. The diffusion coefficient was calculated by fitting the recovery curve using the half-time of recovery ^41,42^. Timepoint zero, *t* = 0 s is defined as the first measured point after the bleaching pulse. The normalized fluorescence intensity *F*_norm_ was calculated from the fluorescence intensity of the region of interest *F*_roi_, the fluorescence intensity of the reference area *F*_ref_, and the background fluorescence intensity *F*_bkgd_ as

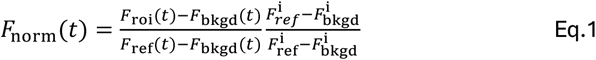

Where *F*^i^ denotes the time average of the fluorescence intensities for *t* < 0 s. *F*_norm_ is subsequently fitted with

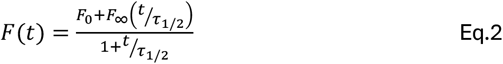

where *F*_0_ is the normalized fluorescence intensity at *t* = 0 s, *F*_∞_ is the fluorescence intensity at *t* → ∞, and τ_1/2_ is the half-time of recovery. From τ_1/2_, the diffusion coefficient *D* can be calculated as

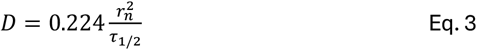

### Scanning probe microscopy measurements

The procedure for measuring and evaluation of rheological properties of condensates with SPM is stated elsewhere in detail ^35^. In short, a 40 µl bulk drop of condensates was prepared on TPP dishes and allowed to settle on the dish for 2 hours. A 5 µl drop from the sample was carefully transferred to the cantilever, to ensure no air bubbles were formed during SPM head placement. SAA-SPH-5UM Si3N4 cantilevers (Bruker, Germany) with a 23 µm height and a 5.16 µm radius hemispheric tip were used. Calibration of the cantilever’s spring constant and amplitude was done using the thermal noise, and the contact-mode method on polystyrene dishes, respectively ^43^. To prevent droplet adhesion, the cantilever was passivated with 1% Pluronic (P2443) for 30 minutes followed by washing with deionized water ^44,45^. After thermal stabilization (10-15 minutes), condensates and cells were indented with a force of 0.3 nN, at a velocity of 1 µm s^−1^ and 40 nm oscillation amplitude. *G*’ and *G*” values were extracted using custom evaluation software (Mathematica 13.2, Wolfram). Condensate radii were measured from phase contrast images in ImageJ. For the hemispheric condensates in buffer the single Hertzian contact model, and for the spheric droplets in medium the double Hertzian contact model were employed ^35^. Condensate experiments were conducted using three independent sample preparations, with 8–16 condensates analyzed per sample. Rheology of cell was performed with four independent sample preparations, with 6–12 cells measured per sample.

To study the interactions between condensates and cells, SV-80 cells, a human fibroblast cell line, were cultured in DMEM supplemented with 10% fetal calf serum, 2.5% HEPES, and 1% penicillin-streptomycin. Once the cells reached 60-70% confluency, the medium was discarded. A 5×5 mm^2^ area was then confined using UV-sterilized adhesive tape (0.12 mm thickness, Grace Bio-Labs, USA) to restrict the condensate application area and minimize the required volume. Thereafter, pK and HS were mixed to a fresh culture medium at 37°C and immediately added on the cells. The cultures were kept at 37°C and 5% CO2 for 2 h, allowing most coalescence events of condensates to cease. Afterwards, the rheological measurements were conducted as stated above on three independent sample preparations, with 12 condensates analyzed per sample.

For adhesion experiments, Arrow™ TL2 cantilevers (NanoWorld, Switzerland) were coated with 10 µg/ml laminin (L2020). Condensates on cells were prepared as stated above and were indented with a force of 1 nN and a velocity of 1 µm s^−1^, and the contact was kept for 60 s before detaching the cantilever at a velocity of 1 µm s^−1^. Experiments were conducted using three independent sample preparations, with 12 condensates analyzed per sample.

### Stacked Hertzian model

Figure 5 schematically depicts the experimental condition for the indenter-condensate-cell stack. The hemispherical Si3N4 probe (subscript 1) is indenting the condensate (subscript 2) which is resting on the cell (subscript 3). While the indenter can be assumed as infinitely rigid, due to the comparable shear moduli of the condensate and the cell, both may deform under the external load by the indenter.

**Figure 5:**
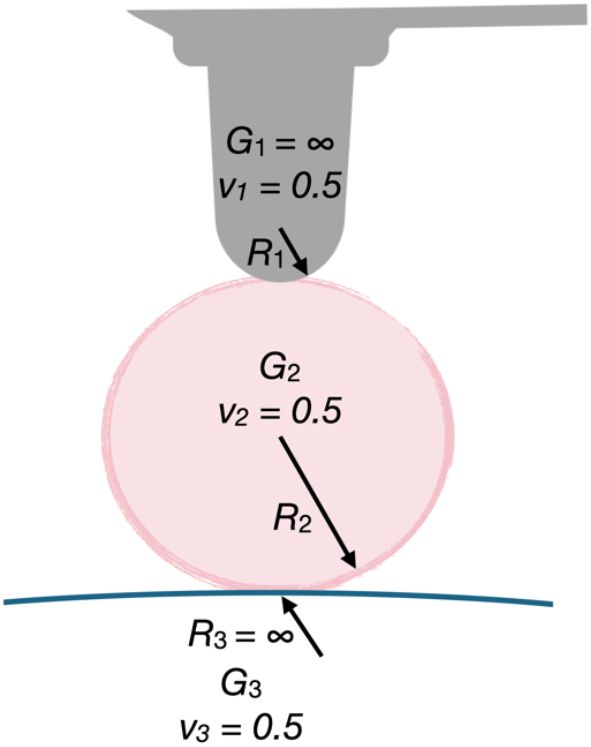
Schematic representation of the rheological stacked Hertzian model and parameters. Subscript 1 : Hemispheric Si_3_N_4_ indenter, *G*_1_ = ∞, *R*_1_ = 5.13 µm, *ν*_*1*_ = 1/2. Subscript 2: Condensate, *G*_2_ is to be determined, *R*_2_ is measured from micrographs, *ν*_*2*_ = 1/2. Subscript 3: Cell, *G*_3_ ~ 1 kPa (exact value is measured separately), *R*_3_ = ∞, *ν*_*3*_ = 1/2.

To calculate the shear moduli of the condensate on the cell, firstly each of the two contacts (indenter-condensate and condensate-cell) can be described by the Hertzian contact model. The contact force between the probe and condensate (denoted as 12) *F*_12_ as a function of deformation at the interface δ_12_ is given by

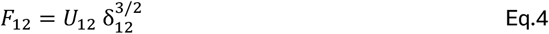

with 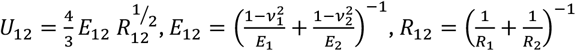, and *E*_1,2_ = 2 *G*_1,2_(2 + ν_1,2_) where *R*_1_ and *R*_2_ are the radii of curvature, *G*_1_ and *G*_2_ are the shear moduli, and ν_1_ and ν_2_ are the Poisson’s ratios of the two bodies, 1 and 2.

Using Hooke’s law, 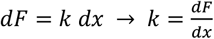, the corresponding spring is given by

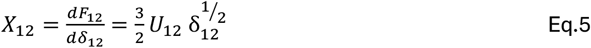

The contact between the condensate and cell (denoted 23) can be described equivalently. Afterwards, the two contacts result in a series connection of two springs (Fig….). In the steady state, the forces at the indenter-condensate interface *F*_12_ and at the condensate-cell interface *F*_22_ are identical, hence *F*_12_ = *F*_22_. With the overall indentation of the cell and condensate (δ = δ_12_+δ_22_), which is known from the SPM head height, the indentations δ_12_ and δ_22_ can be calculated as:

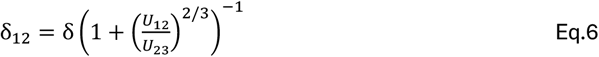

and

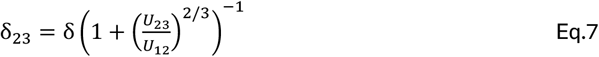

and thus, the respective springs (cf. Eq.5)

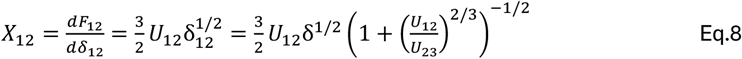

and

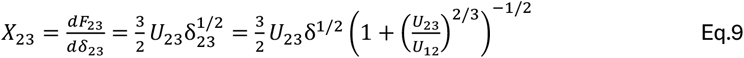

follow. The series connection of *X*_12_ and *X*_22_, *X* is given by

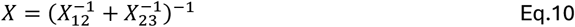

Hence, by separately measuring the shear modulus of the cell (*G*_3_) the shear modulus of the condensate (*G*_2_) can be calculated by means of numerical inversion. It must be noted that *G*_2_ still contains contributions from surface energy at low frequencies which is compensated by subtracting the low-frequency value of *G*′_2_ from *G*_2_(*f*). This is described in detail elsewhere (ref CRPS).

The adhesion forces *F*_ad_ were evaluated with the JPK data evaluation software (version 8.0.168) as the minimum force exerted on the cantilever in the detachment section of the curve. From this value, the surface energy density γ can be calculated as based on the Johnson-Kendall-Roberts (JKR) model ^46^:

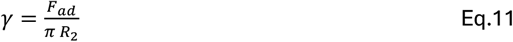

Where R2 is the radius of the condensate.

### Quasi-static approach curve analysis

As demonstrated previously, the surface energy density of the liquid-condensate-interface γ_2_ can be determined from the quasi-static indentation curve of the condensate ^35^. In the present configuration of droplet resting on cell, the indentation curve contains the mechanical response of both the condensate and of the cell. This requires the separation of the deformation of the condensate and the deformation of the cell. To do so, the quasi-static shear modulus of the cell (*G*_2_) needs to be known. This quantity can be determined from the quasi-static indentation curve of the cell by fitting the Hertzian contact model

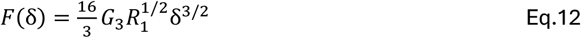

to the experimental data. Knowing *G*_2_, the shear modulus of the condensate *G*_2_ as a function of the indentation δ can be calculated from the experimental force-indentation curve of the condensate resting on the cell by numeric inversion of

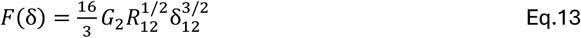

where

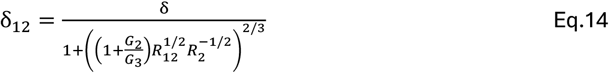

is the indentation at the indenter-condensate interface. As the mechanical response of the condensate is determined by surface energy changes due to spheroidal distortion, *G*_2_ depends on δ_12_ given by

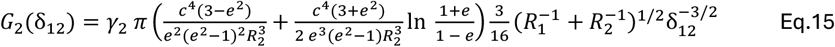

with 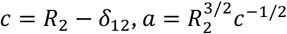, and 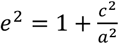.

## Acknowledgments

This project is partly supported by the Dutch Research Council (NWO), Open Competition grants OCENW.XS23.3.105 and OCENW.XS22.4.185. The authors are grateful to the technical support of Laurens Heling and Vahid Sheikhhassani form Medical Systems Biophysics and Bioengineering at Leiden University. The authors also thank Kostas Tassis for assistance with confocal microscopy and FRAP measurements.

## Author Contributions

Conceptualization: A.M., V.S. Methodology: A.M., A.N., O.A. Investigation: A.N. Software: O.A. Visualization: A.N. Formal Analysis: A.N., O.A. Writing & Draft Preparation: A.N., A.M., T.E., Writing-Review & Editing: all authors, Project Administration, Funding acquisition & Supervision: A.M.

## Declaration of Interests

The authors have no conflicting interests to disclose.

## Notes

### Competing Interest Statement

The authors have declared no competing interest.

