## Supplementary Information for "Synthesis and Characterization of Phase-Separated Extracellular Condensates in Interactions with Cells"

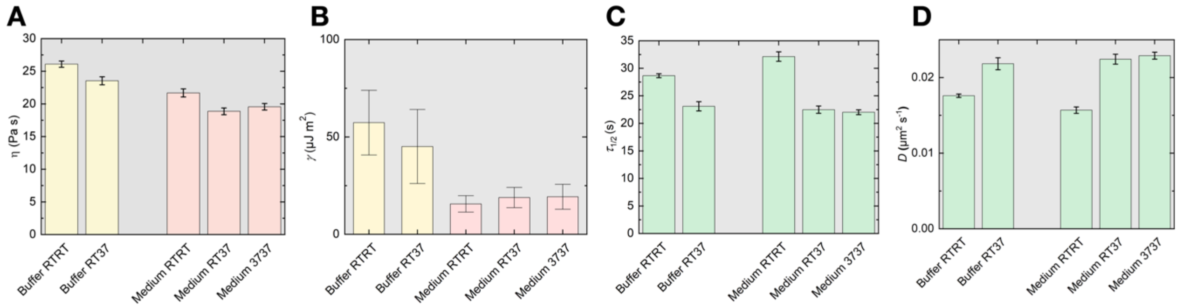


**Supplementary Figure S 1: Comparison of the material properties of pK-HS condensates in various conditions. A)** Viscosity ($\eta$) measured as well as **B)** Surface energy density ($\gamma$) measured with SPM **C)** half-time of recovery ($\tau_{1/2}$) and **D)** diffusion coefficient ($D$) measured with FRAP. **Buffer RTRT:** pK-HS condensates formed in 0.15 M KCl buffer and RT, measured at RT. **Buffer RT37:** pK-HS condensates formed in 0.15 M KCl buffer and RT, measured at 37°C. **Medium RTRT:** pK-HS condensates formed in culture medium and RT, measured at RT. **Medium RT37:** pK-HS condensates formed in culture medium and RT, measured at 37°C. **Medium RTRT:** pK-HS condensates formed in culture medium and 37°C, measured at 37°C.

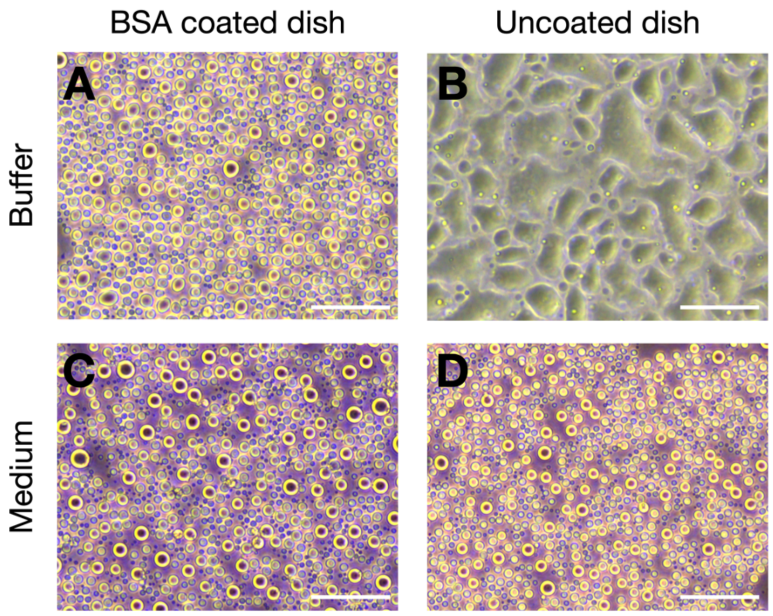


**Supplementary Figure S 2: Comparison of the morphologies of pK-Hs condensates depending on surface passivation and medium. A)** Condensates formed in 0.15 M KCl buffer on BSA coated dish **B)** Condensates formed in 0.15 M KCl buffer on uncoated dish **C)** Condensates formed in culture medium on BSA coated dish **D)** Condensates formed in culture medium on uncoated dish. Scale bar is 50 µm.

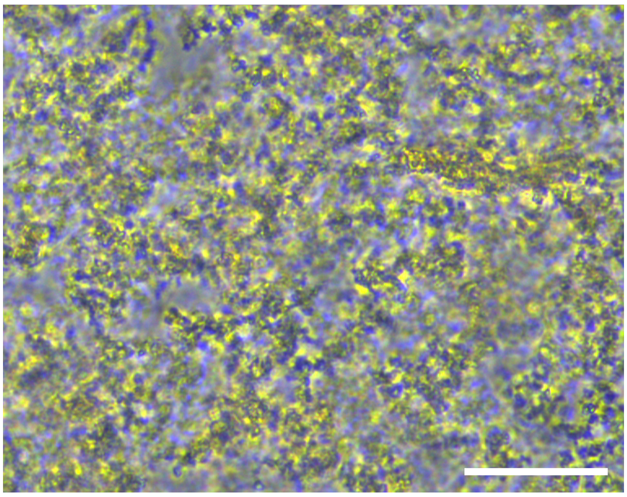

**Supplementary Figure S 3:** pK-H interactions in culture medium. Scale bar is 50 µm.

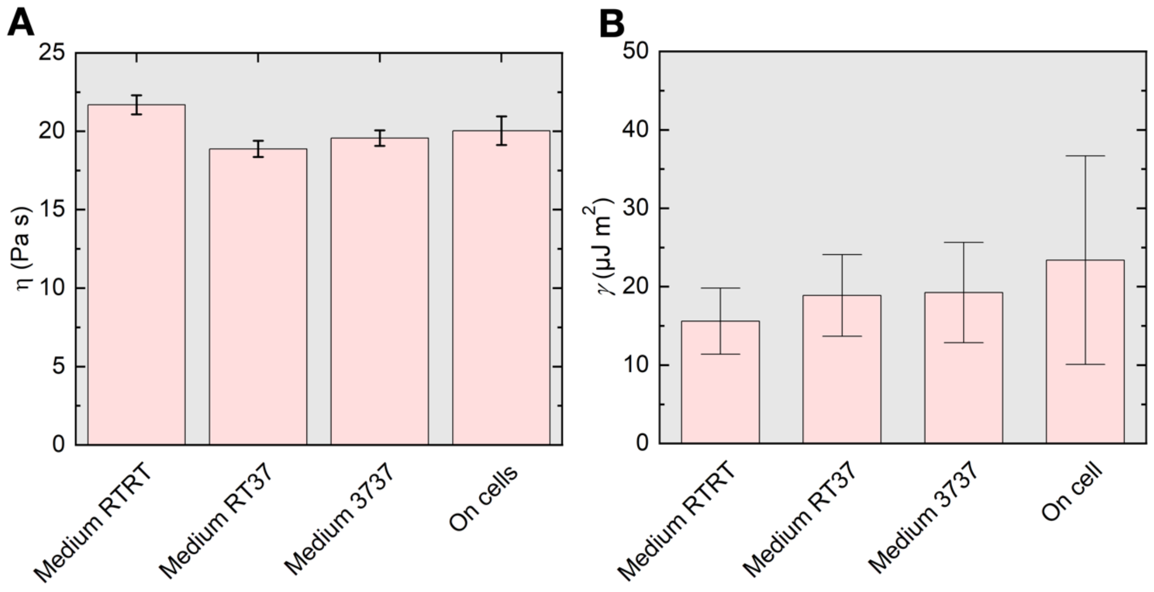


**Supplementary Figure S 4: Comparison of the material properties of pK-HS condensates in various conditions and on fibroblast cells. A)** Viscosity ($\eta$) as well as **B)** Surface energy density ($\gamma$)measured with SPM (Eq. 12-15) for **Medium RTRT:** pK-HS condensates formed in culture medium and RT, measured at RT. **Medium RT37:** pK-HS condensates formed in culture medium and RT, measured at 37°C. **Medium RTRT:** pK-HS condensates formed in culture medium and 37°C, measured at 37°C. **On cells:** pK-HS condensates formed in culture medium and 37°C, measured at 37°C while resting on cells.

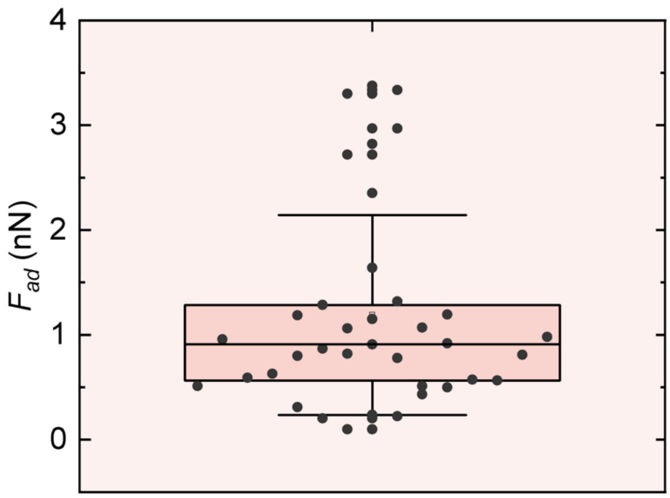


**Supplementary Figure S 5: The distribution of the adhesion forces ($F_{ad}$) between the laminin coated cantilever and pK-HS condensates.**
